# Spotlight on the monitoring of the invasion of a carabid beetle on an oceanic islandover a 100 year period

**DOI:** 10.1101/837005

**Authors:** M. Lebouvier, P. Lambret, A. Garnier, Y. Frenot, P. Vernon, D. Renault

## Abstract

The flightless beetle *Merizodus soledadinus*, originating from the Falkland Islands, was introduced to the sub-Antarctic Kerguelen Islands. We compiled the existing information on ship visits and landings on these islands to confirm the introduction date of *M. soledadinus*. Using data available in the literature, in addition to collecting more than 2000 presence/absence records of *M. soledadinus* over the 1991-2018 period, we tracked changes of its abundance and geographical distribution. The range expansion of this nonflying insect was initially slow, but has accelerated over the past two decades in parallel to local increased abundances of the insect’s populations. Human activities may have facilitated colonization of some localities by *M. soledadinus* which is now widely present in the eastern part of the Kerguelen archipelago. This predatory insect represents a major threat for the native invertebrate fauna; in particular, the wingless flies *Anatalanta aptera* and *Calycopteryx moseleyi* which are locally displaced and/or eliminated by the beetle. If no control measures, let alone eradication, are practicable, it is essential to limit the transport of this invasive insect along with human activities. Since 2006, the Kerguelen Islands have had the status of a nature reserve, making these results of significant interests for the management of this archipelago, and more generally, emphasizing the importance of long-term biomonitoring programmes for assessing and predicting changes in the distribution of invasive organisms. Strict biosecurity measures have now been established at the Kerguelen Islands, with even greater attention paid to visits to remote sites not yet colonized by *M. soledadinus*.

## 1. Introduction

The contribution of anthropogenic activities to biological invasions escalated rapidly over the past decades so that the introduction and spread of non-native organisms have become a global and growing ecological and conservation theme (Parker et al., 1999; Levine and D’Antonio, 2003; Gurevitch and Padilla, 2004). Biological invasions, hereafter referring to the proliferation and impacts of non-native populations introduced through human activities (Richardson et al., 2011) can briefly be sequenced into a six-step continuum: (1) sampling of the specimens in their native geographical area, (2) transport, (3) introduction, (4) establishment, *i.e*. specimens successfully achieve their life cycles, (5) proliferation at the introduction site(s) and (6) spread, *i.e*. geographic expansion (Blackburn et al., 2011; Chabrerie et al. 2019; Simberloff and Rejmanek, 2011). The last two steps of the aforementioned invasion process are now well documented for a range of taxa with for instance studies examining how and why alien individuals have breached the environmental barriers to spread (Thuillier et al., 2006, 2012; and reviewed by Renault et al., 2018 for insects and arachnids) in parallel to works assessing the level of invasiveness of the organisms or invasibility of (micro)habitats (Alpert et al., 2000; David et al., 2017; Hui et al., 2016). Meanwhile, empirical studies that document the early stages of an invasion process are still scarce (Kolar and Lodge, 2001; Jeschke and Strayer, 2005).

In most cases of inadvertent introductions, historical information is not available, and non-native organisms are only first observed when their population densities strongly grow up (Costello and Sollow, 2003; Crooks, 2005) and/or when these specimens start to have economic impacts. This is particularly true for under sampled habitats or cryptic organism populations such as those of many invertebrates because of their small size. Obviously, the ‘*early stage subdetectability*’ can be of considerable importance for authors considering lag effects during invasions (Carey, 1996; Crooks, 2005). There are a few cases where the arrival of alien invertebrates has been detected and monitored, including the invasion process of the ladybird *Harmonia axyridis* (Coleoptera: Coccinellidae) and of the wasp *Vespula velutina* (e.g. Roy and Wajnberg, 2008; Perrard et al., 2009). However, the multiple introduction sites, the multiple introduction events at a single site (Brown et al., 2011), the invasive bridgehead effect (Lombaert et al., 2010), and the overdeveloped anthropogenic activity represent confounding factors that most often prohibit examining the biological and environmental drivers of each step of invasion as separate entities. Importantly, after reading the existing literature, we found no case study of insect invasion that simultaneously documented the four following points: (i) initial site of introduction, (ii) geographical origin of the alien specimens, (iii) way and date of introduction, and (iv) natural spread of the alien species within the area of introduction.

Given the difficulty monitoring the movements of organisms across large geographical areas, estimates of the missing pieces of information – including dispersal rates, geographical distributions, and, more recently, sources of invasion (geographic profiling) – are frequently obtained and mapped using an array of mathematical models (Barbet-Massin et al., 2018; Lustig et al., 2017; Sofaer et al., 2019; Stevenson et al., 2012), and mapping tools are becoming more and more sophisticated Lefcheck 2016). The reliability of these models can be significantly increased when they are calibrated with field observations of the spatial and temporal dynamics of invasion. In particular, presence-absence occurrences over spatio-temporal scales can significantly improve the robustness of models forecasting the expansion of non-native organisms (Schliep et al., 2018). Yet, long-term observations reporting the fine-scale spatial distribution of non-native populations are, to date, rare. When implemented, these records should also consider the use of standard metrics to quantify the invasion level to better forecast invasibility of other habitats/ecosystems and construct efficient invasion indexes to assist managers (Catford et al., 2012).

Several reasons make oceanic islands valuable candidates to conduct invasion ecology studies. These may include *inter alia*, their geographical isolation that limits the influxes of populations from the continents, and their limited terrestrial areas that enable the geographical survey of non-native organisms. Among these oceanic islands, those located in the Southern Ocean are of particular interest especially because numerous multidisciplinary scientific research studies have been carried out over the past decades. On several of these islands, the description and distribution of non-native flora and fauna are documented (see reviews by Frenot et al., 2005; Lebouvier et al. 2011; Hullé et al., 2018) while introduction events are unequally reported (Convey et al., 2011). The absence of extensive anthropogenic activities (no urbanization, industry, agriculture or large environmental pollutions) make sub-Antarctic islands fruitful model systems for invasion science. For instance, several insect species have been introduced and subsequently spread to the Kerguelen Islands (Schermann-Legionnet et al., 2007; Lebouvier et al., 2011) including the carabid beetle *Merizodus soledadinus* (Coleoptera: Carabidae). This insect was introduced to a single site on the Kerguelen Islands (Jeannel, 1940), and by using data available in the existing literature and observations from a general monitoring of invertebrates on the Kerguelen Islands over several decades, it is possible to track changes of its abundance and geographical distribution. The predatory nature of this non-native insect represents a major threat for the native invertebrate fauna from the Kerguelen Islands. In particular, the wingless native flies, *Anatalanta aptera* (Diptera: Sphaeroceridae) and *Calycopteryx moseleyi* (Diptera: Micropezidae), which have very few native predators/competitors (e.g. the rove beetle *Antarctophytosus atriceps*, the spiders *Myro kerguelensis* and *Neomaso antarcticus*), disappeared in some sites colonized by *M. Soledadinus* (Chevrier et al., 1997; Lebouvier et al., 2011).

In the present study, we compiled the existing information on ship visits and landings on the Kerguelen Islands to confirm the introduction date of *M. soledadinus*. Then, a 105-year time series of the spreading process of *M. soledadinus* is reported, starting from its introduction to present day. The relative abundance, seasonal phenology of this ground beetle, and its impact on two native species from the Kerguelen Islands, the wingless flies *A. aptera* and *C. moseleyi*, are also reported.

Such a long-term data set is rare and very useful in the field of biological invasions, but our results are also valuable in terms of management. Since 2006, the Kerguelen Islands have had the status of a nature reserve. The manager of this reserve is directly interested in information on the distribution and dispersion of the invasive beetle for the implementation of biosecurity measures, particularly when travelling to areas not yet colonized by *M. soledadinus*.

## 2. Material and methods

### 2.1. Study area

The present study was conducted on the Kerguelen Islands (48°30’-50°S, 68°27’-70°35’E), a sub-Antarctic archipelago located in the Southern Indian Ocean more than 3500 km away from the African and Australian coasts. This wide archipelago (7200 km^2^) consists of a main island (6500 km^2^), ten or so small islands (1–200 km^2^), and numerous islets. The highest point is Mont Ross (1850 m), and an ice cap (Glacier Cook) is present in the western sector (Fig. 1). There is no permanent population, but the research station (Port-aux-Français), established in 1950, hosts 50–100 persons all year round. Gravel roads are restricted to the vicinity of the research station, and human activities, i.e. research, logistics and –at very low level– tourism, mainly concern the eastern sector of the archipelago (Péninsule Courbet, Golfe du Morbihan, Péninsule Jeanne d’Arc). Visits to more remote sites are limited because they involve the use of vessels or helicopters that are not permanently available. In 2006, these islands were given the status of Nature Reserve, the highest protection available under French law, and some wilderness areas were classified as “strict nature reserve” where human access, use, and impacts are strictly controlled and limited.

**Fig. 1.**
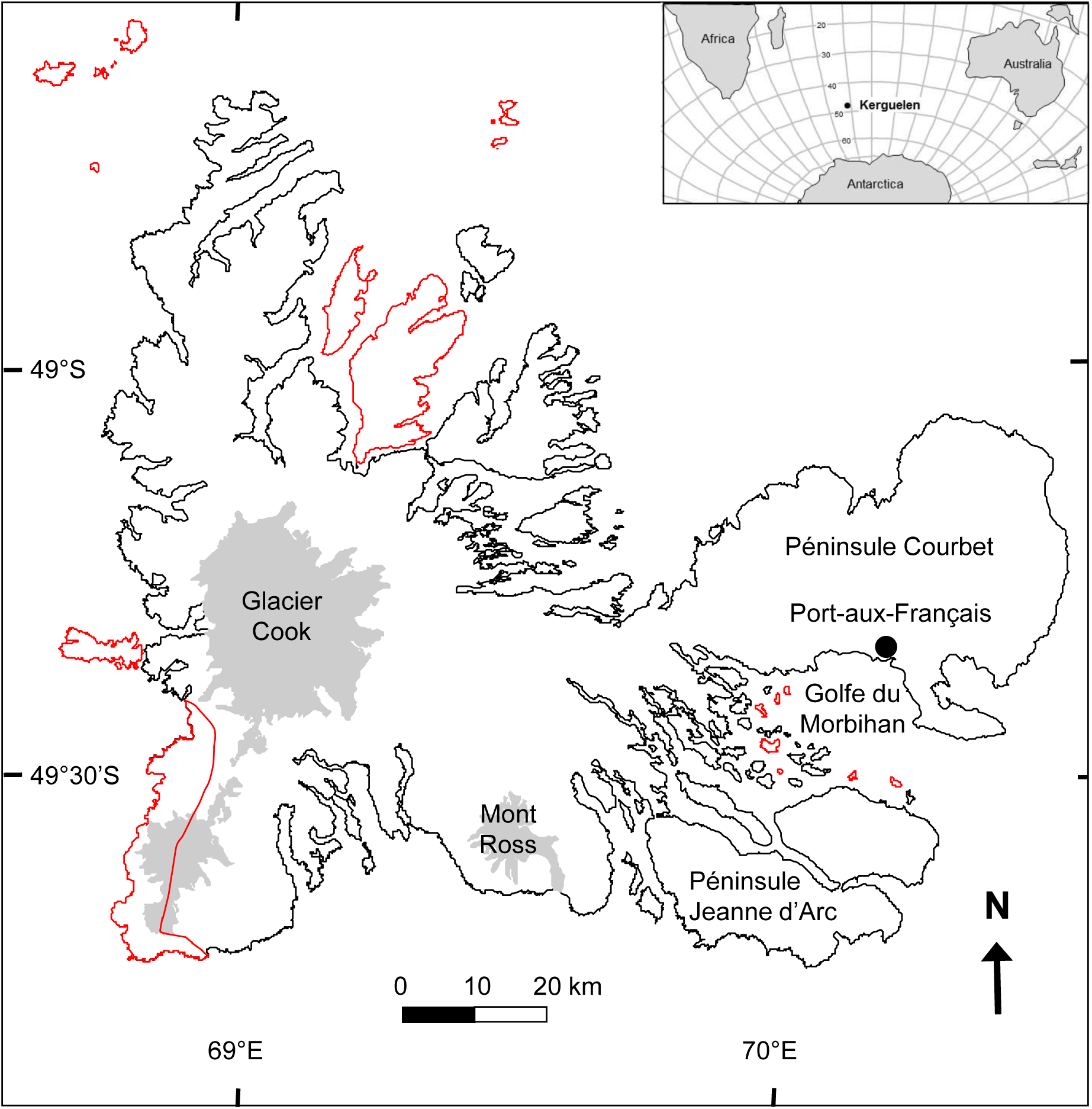
Location of the Kerguelen Islands in the Southern Hemisphere and map of the archipelago. All islands are part of a Nature Reserve. Wilderness areas classified as “strict nature reserve” are indicated in red.

### 2.2. Biological models

#### Merizodus soledadinus

Guérin-Méneville 1830 (Coleoptera, Carabidae) is a flightless carabid beetle which originates from southern South America (Patagonia) and the Falkland Islands (Darlington, 1970; Johns, 1974; Robinson, 1984; Niemelä, 1990; Casagranda et al. 2009). It was described (as *Trechus soledadinus*) by Guérin-Méneville, from the Falkland Islands (Soledad Bay), in 1830. Later, Enderlein (1912) also reported the insect from the Falkland Islands as *Dormeyeria soledadina*. In 1940, Jeannel named the species *M. soledadinus* which turned to *Oopterus soledadinus* by Johns (1974). But Lalouette (2009) has restored *M. soledadinus*, a denomination further confirmed by Voisin et al., (2017). *Merizodus soledadinus* has been accidentally introduced to the Kerguelen Islands (first record in 1939, Jeannel, 1940) and to South Georgia, 54°S (first record in 1963, Darlington, 1970). Adults were described as being active at night (Ottesen, 1990) and found during the day beneath stones and kelp belts (Ernsting, 1993; Renault et al., 2015).

#### Anatalanta aptera

Eaton 1875 (Diptera, Sphaeroceridae) is a sub-Antarctic wingless fly that is found on the Crozet, Kerguelen, Heard and McDonald Islands. On the Kerguelen Islands, this species is present from sea level to more than 600m a.s.l. and is active all year round. Larvae and adults are saprophagous and feed on decayed organic matter. *Anatalanta aptera* is abundant in many habitats, especially in seabird colonies, under carrions and on coastal areas enriched by seaweeds (Vernon, 1981; Chevrier et al., 1997).

#### Calycopteryx moseleyi

Eaton 1875 (Diptera, Micropezidae) is a sub-Antarctic wingless fly that is found on the Kerguelen, Heard and McDonald Islands. Its larvae feed preferentially on the Kerguelen cabbage *Pringlea antiscorbutica*, but can also frequently be found under decaying seaweeds along the seashore and on decayed organic matter in penguin rookeries (Tréhen et al., 1987).

### 2.3. Spatio-temporal monitoring of the distribution and abundance

#### 2.3.1. Date of introduction and spread

Within the existing literature, the supposed introduction date of *M. Soledadinus* to the Kerguelen Islands varies from 1800 (Jeannel, 1940) to 1913 (Chevrier et al., 1997) or to 1927 (Jeannel, 1962). We first reviewed the available literature to clarify the introduction route and date of this insect and its subsequent geographical distribution throughout the Kerguelen Islands (Jeannel, 1940, 1964; Tréhen and Voisin, 1984; Briot, 1990, Dreux et al., 1992; Chevrier, 1996; Arnaud and Beurois, 1996; Delépine 2002). Then, to assess the changes in the geographic distribution of the beetle, we analyzed more than 2000 presence/absence records of *M. soledadinus* collected over the 1991-2018 period. The records were projected in a grid cell of 1km for the drawing of the distribution maps of the insects.

#### 2.3.2. Quantifying the invasion level of Merizodus soledadinus

Since 2005, a systematic survey of the geographical distribution and abundance of *M. soledadinus* has been carried out, mostly along the coastline where this alien insect was confined in the early stages of its introduction and spread throughout the Kerguelen archipelago (Chevrier et al., 1997). For each prospected site, we noted the GPS coordinates of the record and the occurrence (presence/absence, index of abundance) of *M. soledadinus* and of the two wingless native flies. The nature of micro-habitats hosting the three insect species was also recorded: stranded seaweed, stones, carrions, and leaves of the Kerguelen cabbage. Inland sites were also prospected by conducting observations on transects perpendicular to the shoreline and on altitudinal transects. To refine *M. soledadinus* distribution limits at the edge of colonized areas, observations were done ca. every 100 meters on flat transects and ca. every 20m in elevation for altitudinal transects. Observations were stopped when no *M. soledadinus* were observed at two consecutive observation points. Preliminary tests showed that a 10-min search by one person was appropriate to detect the presence and estimate the abundances of the three insect species, even in the case of low abundance levels. A semi-quantitative index was designed according to the number of adults found during the 10-min search: (0) absence, (1) low abundance, 1-30 adults, (2) medium abundance, 31-100 adults, and (3) high abundance, more than 100 adults.

### 2.4. Seasonal phenology and assessment of the ecological impact of the predatory carabid beetle on native entomofauna

To specify the seasonal activity and abundance of *M. soledadinus, A. aptera* and *C. moseleyi*, we used data from a multi-year invertebrate monitoring program conducted at several sites on the Kerguelen Islands using pitfall traps and yellow traps.

Using the abundance index recorded during the geographical survey conducted from December 2004 to March 2006, a first assessment of the ecological impact of *M. soledadinus* on the two native flies *A. aptera* and *C. moseleyi* was conducted and took into account the period of activity of these three insect species. We focused on the four following binomial comparisons: (i) *M. soledadinus* vs. *A. aptera*: seashore, under seaweeds, stones… (338 records), (ii) *M. soledadinus* vs. *A. aptera*: inland, i.e. at more than 50m from the seashore, under stones and carrions (379 records), (iii) *M. soledadinus* vs. *A. aptera*: under carrions, along the seashore and inland (155 records), (iv) *M. soledadinus* vs. *C. moseleyi*: seashore, under seaweeds and stones (177 records).

### 2.5. Statistical analyses

GIS tools (ArcGIS 10.4, Esri) were used to map the changes in the geographical distribution of *M. soledadinus* over time. The frequency distribution of the abundances of *M. soledadinus, A. aptera*, and *C. moseleyi* were represented in contingency tables which represent marginal (sum of each column, sum of each line) and grand (total number of individuals) totals. Expected frequencies were first computed from the totals assuming that there were no relationships among cells which would result in similar values between expected and observed frequencies. To assess differences among proportions, Chi-square tests were conducted (whenever classes with low frequencies occurred, frequencies from adjacent classes were pooled) as well as Fisher’s exact test (when classes pooling resulted in a 2×2 table). The analyses were conducted with Minitab 13 (Minitab Inc., StateCollege, PA.).

## 3. Results

### 3.1. Spatio-temporal monitoring of the geographical distribution of Merizodus soledadinus

*Merizodus soledadinus* was first observed on the Kerguelen Islands in February 1939 by Jeannel (1940) who then actively searched for this insect in various locations within the archipelago. Jeannel reported the presence of *M. soledadinus* (more than one thousand individuals) only in the surroundings of the remnant farm buildings of Port-Couvreux (Fig. 2). He concluded that the restricted geographical range of *M. soledadinus* attested to a recent introduction. He first hypothesized the activities of American sealers as the introduction source of *M. soledadinus* (the ship ‘*Hillsborough*’ landed at Port-Couvreux in 1799). However, he later revised his statement and estimated that *M. soledadinus* would have been found in a larger area of the Kerguelen Islands if introduced around 1800 (Jeannel, 1964). Thus, he then proposed that *M. soledadinus* individuals were introduced to the Kerguelen Islands when the buildings (piggery, sheepfold) of Port-Couvreux were enlarged in 1927-1928 (Delépine, 2002). Yet, this assumption can be refuted, as the attempt to start sheep farming in 1927 was conducted with animals loaded during a call in Durban (South Africa) by the ship ‘*Lozère*’ conveying material from Le Havre (France) to the Kerguelen Islands. Arnaud and Beurois (1996) listed all of the ship landings that occurred at Port-Couvreux, and from this list, we identified the ‘*Jacques*’, a vessel belonging to René Bossière, as the most serious source of the introduction of *M. soledadinus*. The ship left Swansea (Wales) in February 1913, sailed (via Montevideo, Uruguay) to the Falkland Islands where it stayed about one month and loaded equipment and ca. 1600 sheep. Tussock grass was used as fodder to feed sheep during their transport to the Kerguelen Islands and most probably contained specimens of *M. soledadinus*. In August 1913, the ‘*Jacques*’ arrived at Port-Couvreux where the 1150 surviving sheep were unloaded (Arnaud and Beurois, 1996). An additional argument in favour of the introduction of *M. soledadinus* via the unloading of sheep is the presence at Port-Couvreux (and to date nowhere else on the Kerguelen Islands) of *Trisetum spicatum* (L.) K. Richt., a grass species from cold regions of the Northern and Southern hemispheres which is also present on the Falkland Islands (Frenot et al., 2001).

**Fig. 2.**
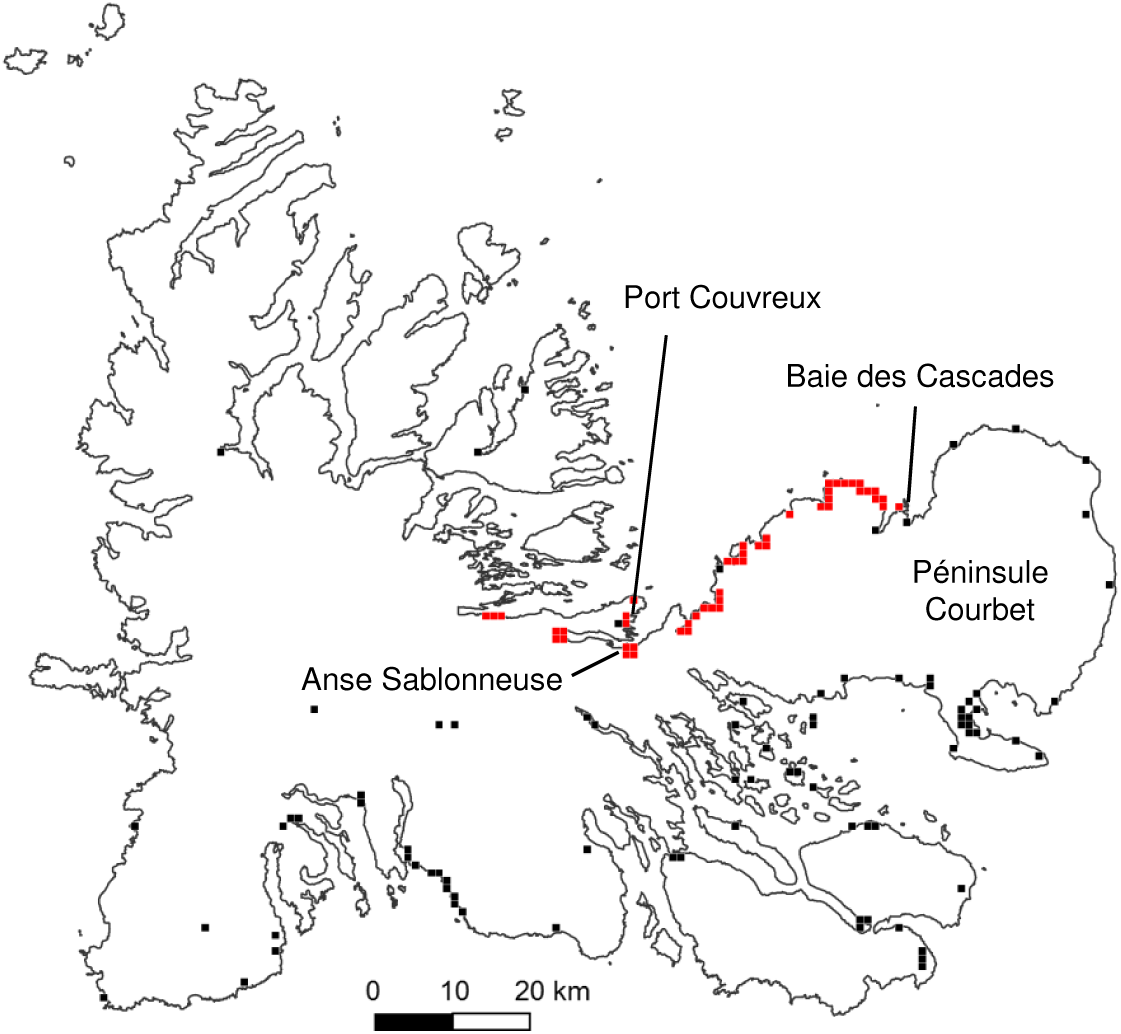
Observations on the distribution of *Merizodus soledadinus* on the Kerguelen Islands in 1983 (after Dreux et al., 1992). Observations are plotted on a one kilometer grid; red square = presence, black square = absence.

The flightless carabid beetle has long remained restricted to the vicinity of its initial introduction site at Port-Couvreux (Jeannel, 1940). Then, it gradually colonized coastal habitats along the northeast coast of the Péninsule Courbet (Fig. 2) up to the Baie des Cascades in 1983 (Dreux et al., 1992). In the early 1990s (Chevrier, 1996), *M. soledadinus* expanded further on the Péninsule Courbet and reached Cap Cotter (Fig. 3). It also colonized an island in front of Port-Couvreux (Ile du Port), new locations remote from its original point of introduction (Port-Phonolite and Port-Jeanne d’Arc, an ancient whaling station where scientists, but also tourists, regularly landed), and one island of the the Golfe du Morbihan (Ile Haute) (Fig. 3). Subsequently, the colonization process accelerated considerably, as shown by the surveys conducted between 2005 and 2007 (Fig. 4). Numerous sites far from Port-Couvreux were colonized. In some instances, it has been possible to have accurate estimates of the date of establishment of populations of *M. soledadinus* through regular trapping sessions set up in the mid-1990s for long-term monitoring of invertebrates. At the research station (Port-aux-Français), the first collections of *M. soledadinus* occurred in June 2000, and they became very regular from 2002 onwards. In the Golfe du Morbihan, the carabid beetle was trapped for the first time in 1997 on Ile Guillou and in 2002 on Ile aux Cochons. In 2007, the insect had invaded almost the entire coastline around the Golfe du Morbihan and many islands in this bay, including islets that have rarely been visited. Specimens of *M. soledadinus* were also reported from several additional sites: Port Fleuriais (north of Port-Couvreux) and, on the east coast, in the vicinity of an isolated hut (Estacade). Of note, *M. soledadinus* was recorded inland in the vegetation along rivers (e.g. Gave de l’Azorella) but also on fell-field in altitude (Plateau du Larzac, in 2005, at 290 m asl; Plateau Central, in 2006, five records between 278 and 358 m asl). In 2018 (Fig. 5), the distribution of the insect significantly widened in the interior of the Péninsule Courbet particularly in the valley between Port-aux-Français and Port Elizabeth. On the east coast, *M. soledadinus* colonized the entire Baie Norvégienne and has now been recorded between Estacade and Cap Digby. In the Golfe du Morbihan it was present on all prospected islands and islets. To date, no *M. soledadinus* have been observed in the western part of the Kerguelen Islands (Massif Gallieni, Péninsule Loranchet and Péninsule Rallier du Baty).

**Fig. 3.**
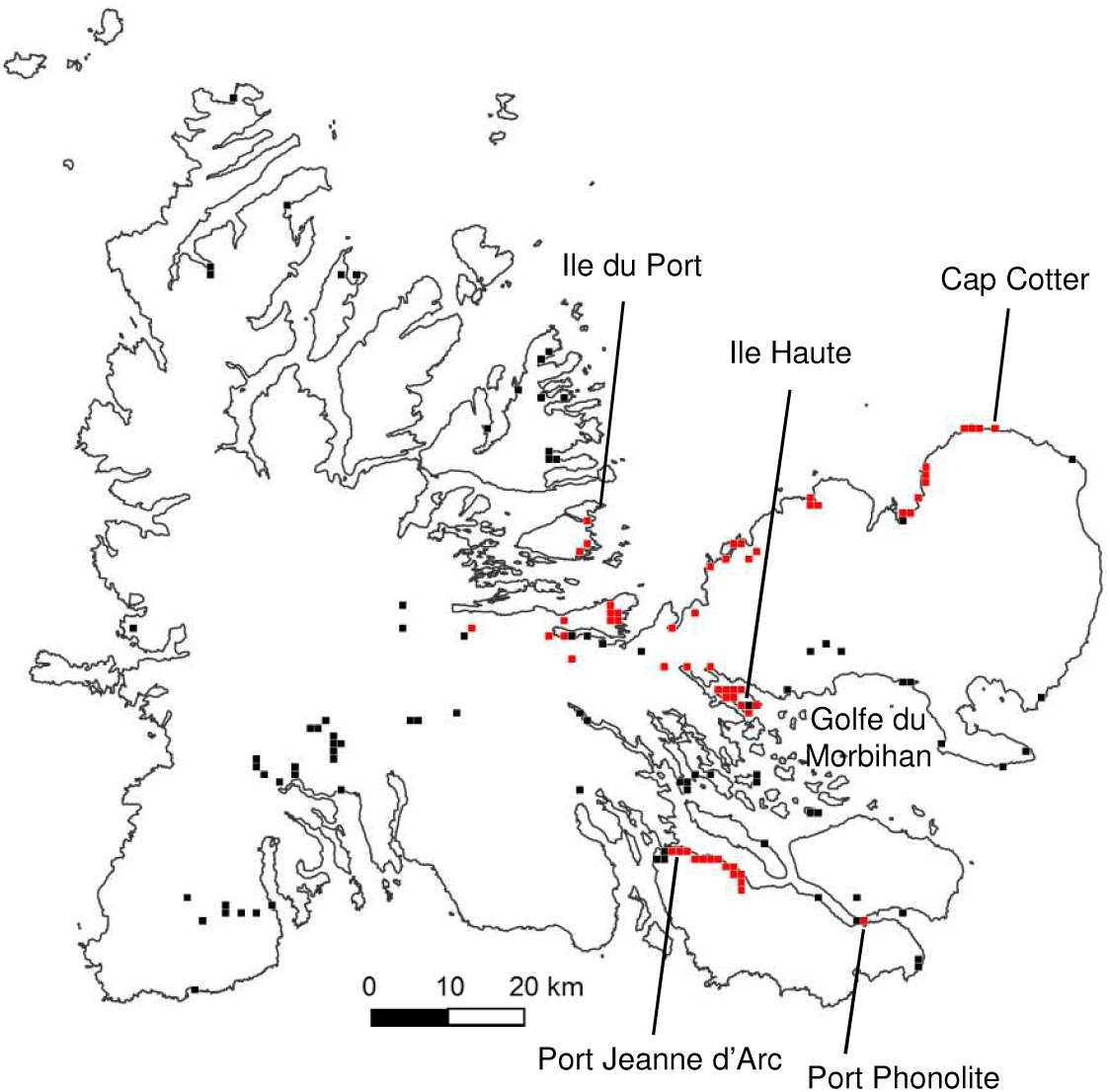
Observations on the distribution of *Merizodus soledadinus* on the Kerguelen Islands between 1991 and 1995 (after Chevrier, 1996). Observations are plotted on a one kilometer grid; red square = presence, black square = absence.

**Fig. 4.**
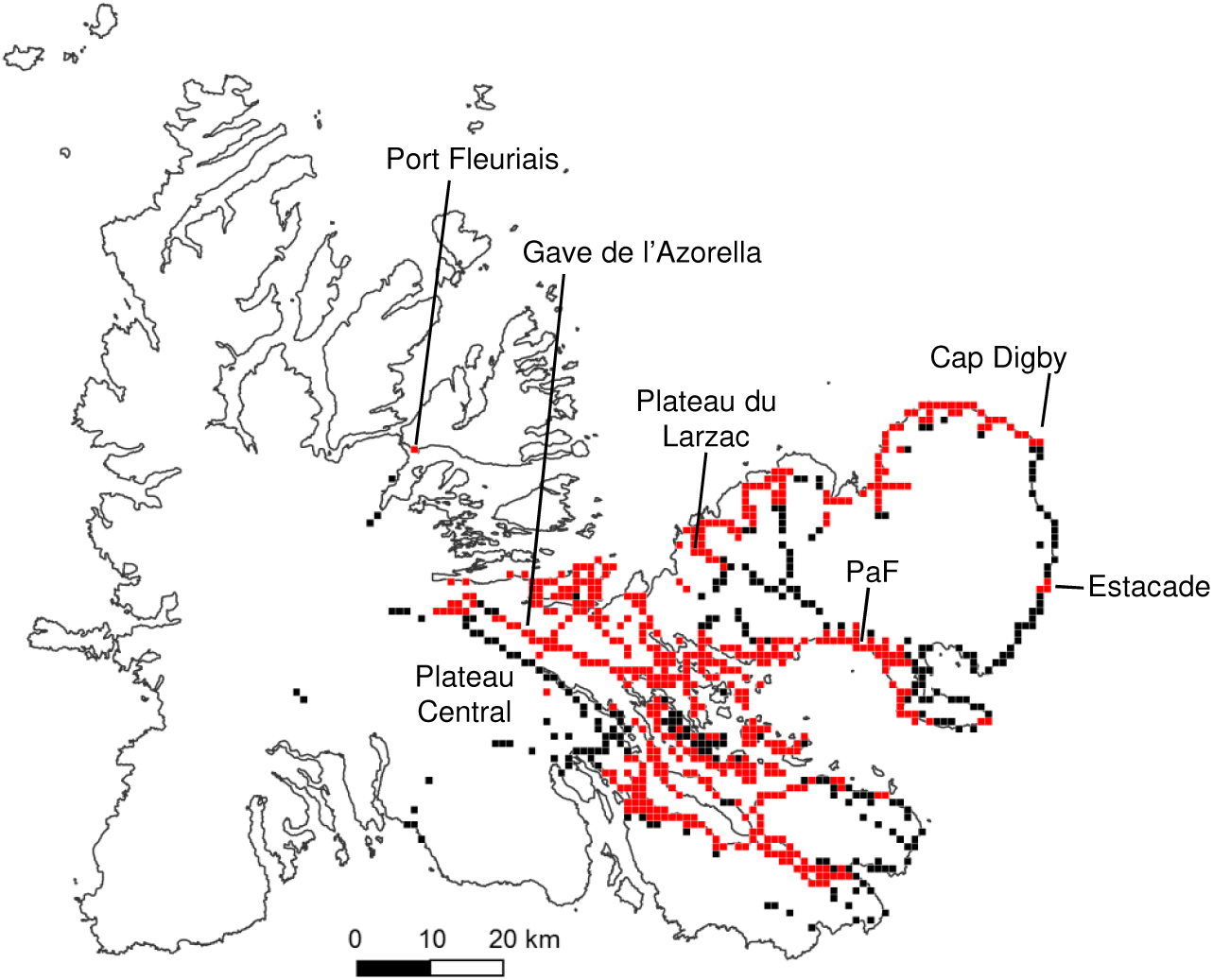
Observations on the distribution of *Merizodus soledadinus* on the Kerguelen Islands between 2005 and 2007. Observations are plotted on a one kilometer grid; red square = presence, black square = absence.

**Fig. 5.**
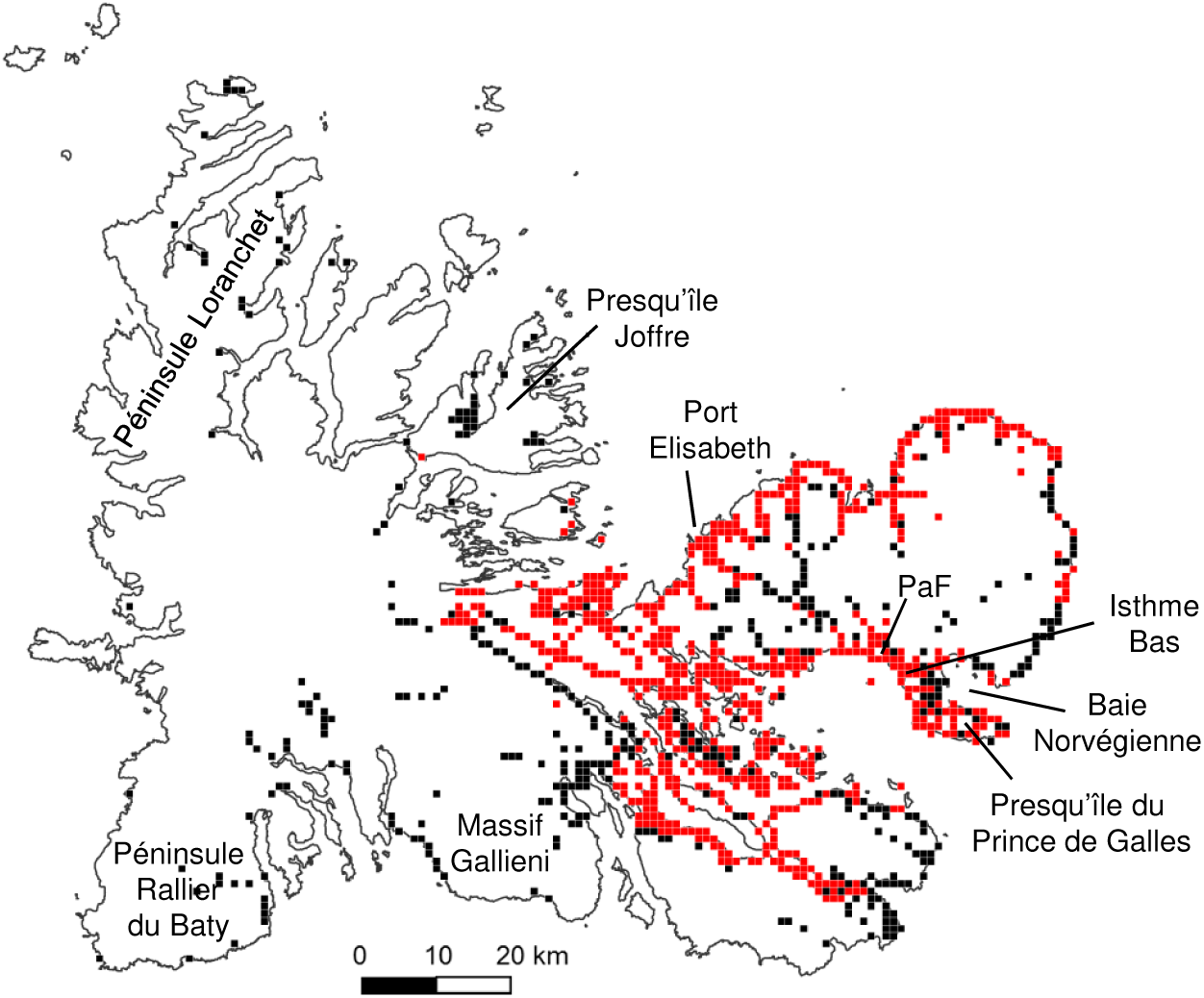
Map of all observations on the distribution of *Merizodus soledadinus* on the Kerguelen Islands between 1939 and 2018. Observations are plotted on a one kilometer grid; for each cell of the grid the status (presence in red or absence in black) corresponds to the most recent observation.

An abundance index of *M. soledadinus* was determined for each of the 1164 locations where the species was recorded between 2005 and 2018 (low abundance, n=680; medium abundance, n=265; high abundance, n=219). The carabid beetle is particularly abundant in the vicinity of Port-aux-Français, on Presqu’île du Prince de Galles, and in between Cap Cotter and Cap Digby. It is interesting to note that high abundances of *M. soledadinus* have been observed throughout the colonized area, on the coast, on the islands, and sometimes inland, both for areas colonized for several decades and for areas recently colonized. So the species can quickly become abundant after colonization. This is also illustrated by the results of monthly trapping that started in 2005 on both sides of Isthme Bas: the number of captured adult *M. soledadinus* on the eastern coast, colonized in 2011, rapidly reached similar levels as those recorded from the west coastcolonized between 2000 and 2005 (Fig. 6).

**Fig. 6.**
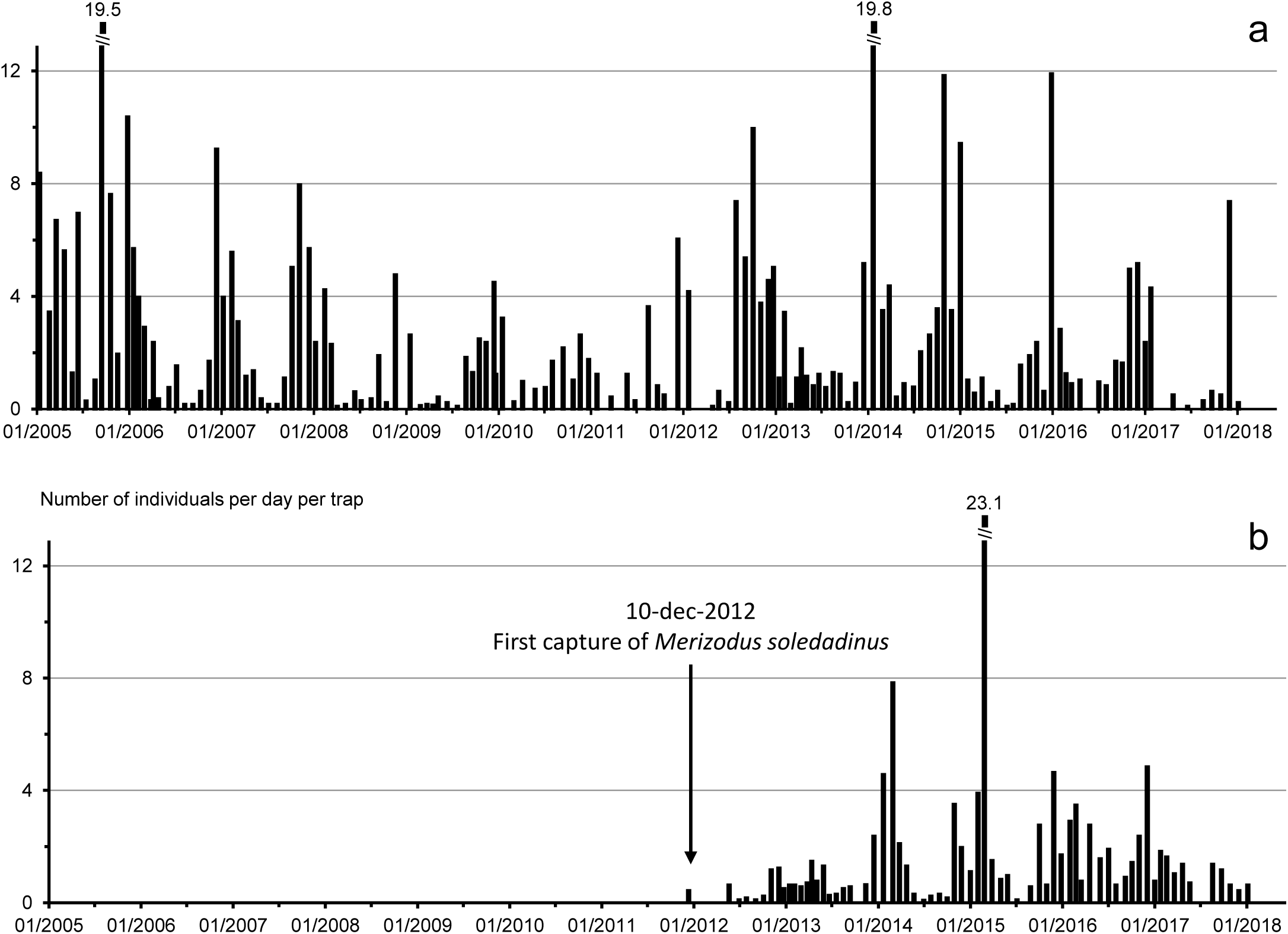
Captures of *Merizodus soledadinus* at two sites between 2005 and 2018 (three pitfall traps open during five days every two or three weeks), a. west coast of Isthme-Bas, site colonized between 2000 and 2005, b. east coast of Isthme-Bas, first observation on this site in 2012.

### 3.2. Impact of Merizodus soledadinus *on the native fly* Anatalanta aptera

During the period of this study, from December 2004 to March 2006, specimens of *M. soledadinus* and *A. aptera* exhibited continuous activity, yet it highly decreased in winter (Fig. 7a, b). Less than 10% of the abundance index taken into account for this analysis (n=862) were recorded in July and August.

**Fig. 7.**
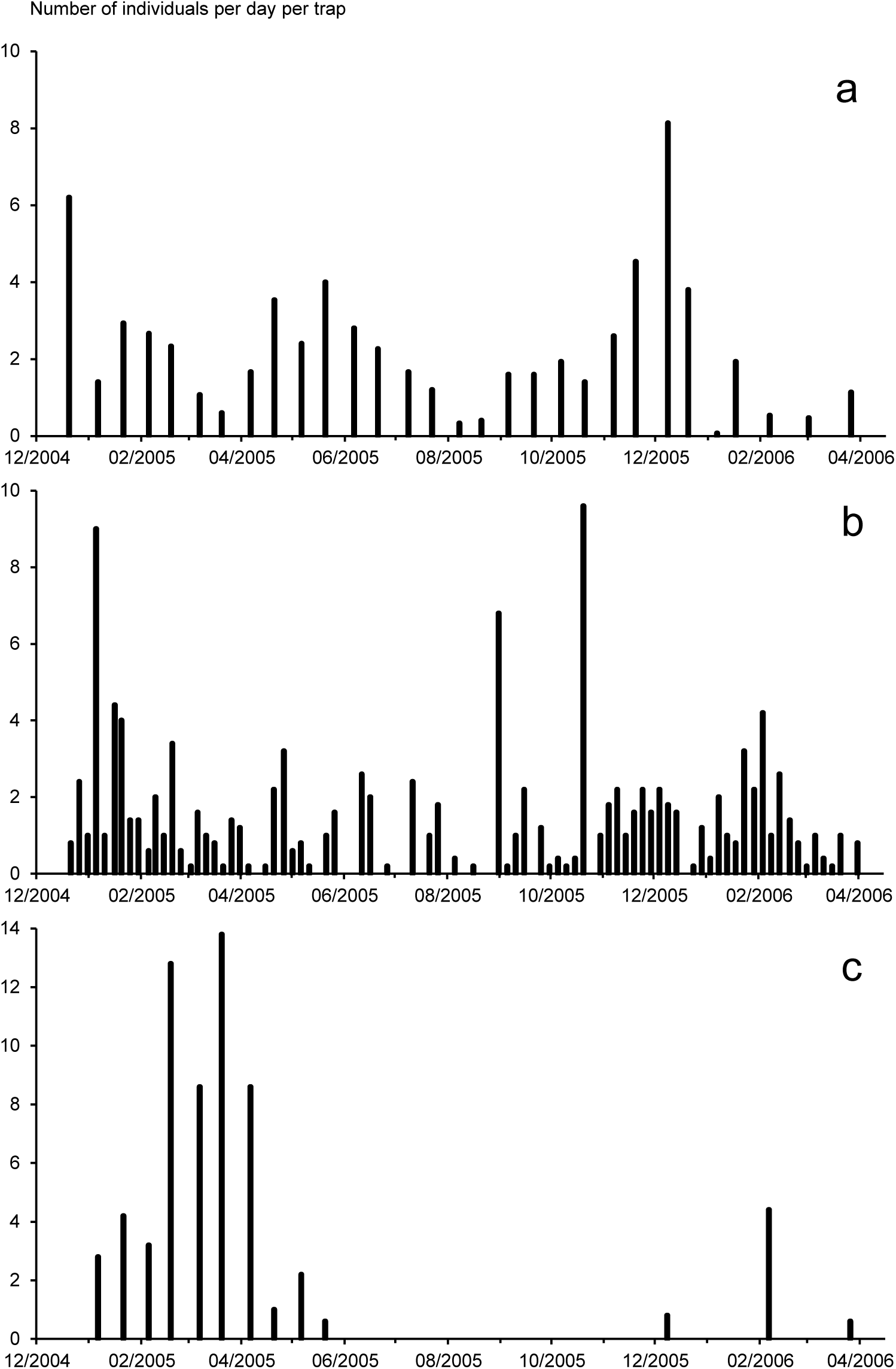
Pattern of activity of three species of invertebrates on the Kerguelen Islands a. *Merizodus soledadinus* (three pitfall traps open during five days every two or three weeks) b. *Anatalanta aptera* (one baited trap continuously open, insects collected every five to ten days c. *Calycopteryx moseleyi* (one yellow trap open during five days every two or three weeks)

The abundance of *A. aptera* (Table 1) was not independent from the abundance of *M. soledadinus* along the seashore (n=338), inland (n=379) or under carrions (n=155): *A. aptera* was more often present and abundant than expected when *M. soledadinus* was absent, and, conversely, the fly was more often than expected absent or at low abundance when *M. soledadinus* was present.

**Table 1.**
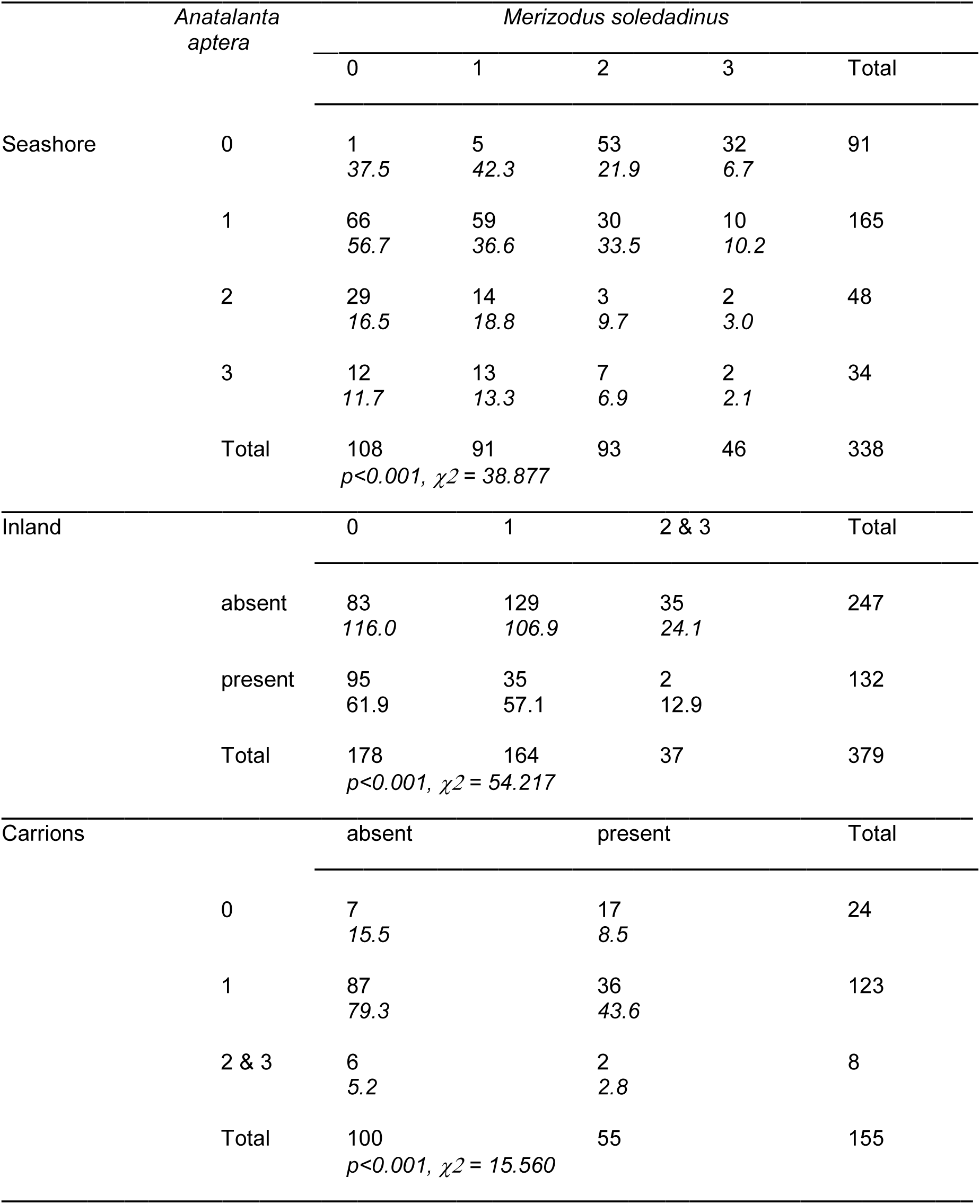
Frequency distribution of the abundances of adult *Anatalanta aptera* and *Merizodus soledadinus* in three habitats. Code for abundances according to the number of adults found during a 10 min active search: 0 = absent; 1 = low abundance, 1-30 adults; 2 = medium abundance, 31-100 adults; 3 = high abundance, more than 100 adults. Expected frequencies in italics.

### 3.3. Impact of Merizodus soledadinus *on the native fly* Calycopteryx moseleyi

Adults of *C. moseleyi* were only trapped from January to March 2005 (Fig. 7c). The abundance index of *M. soledadinus* and *C. moseleyi* taken into account for this analysis were recorded during this period (n=177). The abundances recorded for the two species (Table 2) were not independent (Chi-2= 36.007, p < 0.001). Numbers of *M. soledadinus* negatively affect the abundance of adult *C. moseleyi*: when the ground beetle was present within the habitat, *C. moseleyi* was more often absent (0) and less often abundant (abundance index 2 or 3) than predicted.

**Table 2.** Frequency distribution of the abundances of adult *Calycopteryx moseleyi* and *Merizodus soledadinus*. Code for abundances according to the number of adults found during a 10 min active search: 0 = absent; 1 = low abundance, 1-30 adults; 2 = medium abundance, 31-100 adults; 3 = high abundance, more than 100 adults. Expected frequencies in italics.

|  |  | *Merizodus soledadinus* |  |  |
| --- | --- | --- | --- | --- |
|  |  | absent | present | Total |
| *Calycopteryx moseleyi* | 0 | 15<br>*31.5* | 97<br>*73.5* | 112 |
|  | 1 | 30<br>*16.8* | 26<br>*39.2* | 56 |
|  | 2 & 3 | 6<br>*2.7* | 3<br>*6.3* | 9 |
|  | Total | 51 | 126 | 177 |
|  |  | *p < 0.001, $\chi^2 = 32.925$* |  |  |

Additional information on the impact of *M. soledadinus* on *A. aptera* and *C. moseleyi* was provided by trapping results from a coastal site on Ile Guillou (three pitfall traps opened monthly during five days from January 1994 to July 2003 and then from March 2006 to November 2007). Both flies were present at this site at the arrival of *M. soledadinus* (first trapped in July 1998 on this island). The ground beetle was then regularly trapped until 2003 (between 0.1 and 0.7 individual per trap per day in 13 of the 54 trapping sessions). At the same time, a decrease of the captures of *A. aptera* and *C. moseleyi* was recorded (Fig. 8). When trapping was resumed in 2006, *M. soledadinus* was caught at every trapping session and its abundance was much higher (1 to 11 individuals per trap per day with a maximum value of 46 in December 2006). At the same time the number of *A. aptera* caught was very low and no more *C. moseleyi* was trapped.

**Fig. 8.**
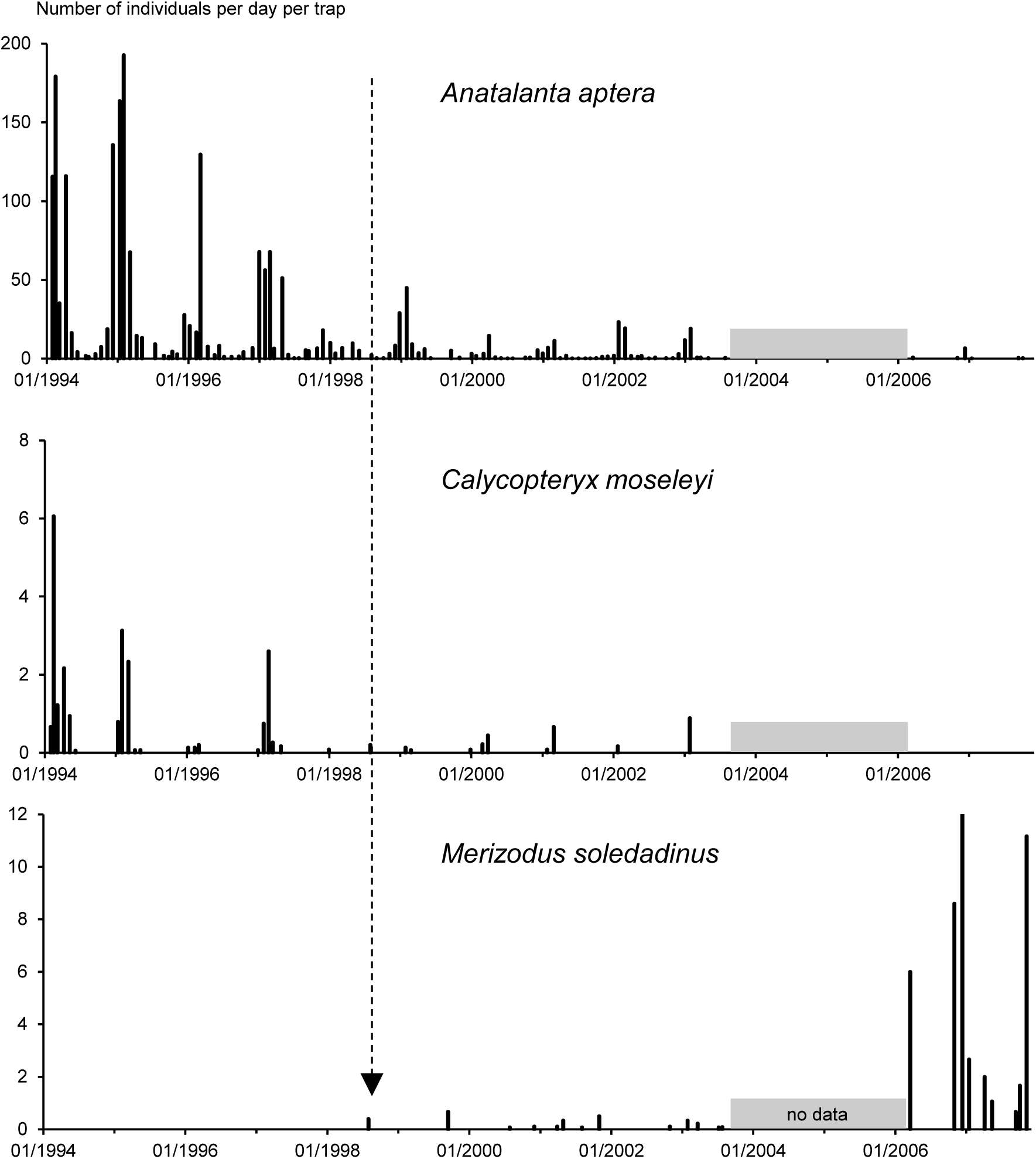
Captures of *Anatalanta aptera, Calycopteryx mosleyi*, and *Merizodus soledadinus* at Ile Guillou between 1994 and 2007 (three pitfall traps open during five days every month). The arrow indicates the date (30/07/1998) of the first capture of *Merizodus soledadinus* on this site.

## 4. Discussion

Geographical spread of non-native organisms represent a hot topic in invasion science and a significant component of the management procedures that can be adopted to mitigate alien-related problems (Hulme et al., 2008). Most often, human-assisted spread of non-native organims can result in multiple introduction points. As a result, the large spatial scales that must be covered impede in-field monitoring of non-native species distribution and abundance (Veldtman et al., 2010). Even if our understanding of range expansion can be implemented through empirical and modeling analyses, the accuracy of these estimates can be significantly improved by incorporating real space-time distribution data (Kadoya et al., 2009).

In the present work, we took advantage of the non-intentionally introduced carabid beetle *M. soledadinus* on the Kerguelen Islands to report a unique space-time in-field monitoring of the geographical spread of a non-native insect. After its initial introduction in 1913 at Port-Couvreux, the insect proliferated on this site over several years, at least until 1939, when Jeannel (1940) actively searched for its presence. Regarding the subsequent expansion of the distribution of *M. soledadinus* in this archipelago, we suggest that Allee effect, which refers to any process whereby any component of individual fitness is correlated with population size (Allee, 1931), could be one explanation for this observed lag phase at the initial stage of the invasion process on the Kerguelen Islands (also see the review of Chabrerie et al., 2019 which lists the different concepts explaining invasion dynamics). With an Allee effect, the introduced species increases very slowly at first, until it builds up a large enough population size at which point there may be a rapid escalation of spread (see Taylor and Hastings, 2005 for a review). Subsequently, from the 1970s onwards, the increase in average temperatures likely accentuated the phenomenon (Lebouvier et al., 2011).

### 4.1. 1983, the start of in-field monitoring of the non-human assisted geographical spread of M. soledadinus on the Kerguelen Islands

Farming (and human presence) stopped in 1932 at Port Couvreux. There was nearly no human presence on the Kerguelen Islands until the early 1950s, when the research station Port-aux-Français was established. In the 50s and 60s, human activities were mainly restricted to the surroundings of the station and the islands of the Golfe du Morbihan. As a result, individuals of *M. soledadinus* spread over the Kerguelen Islands without any human assistance, at least until they reached Port-aux-Français during the 1995-1996 period. In 1983, specimens of *M. soledadinus* were sampled at Anse Sablonneuse, 25 km away from Port-Couvreux by terrestrial means but far closer if we are considering the distance as the crow flies. The observed distribution of *M. soledadinus* on this date (Fig. 2) strongly suggests that some imagoes directly crossed the inlet as the coastal distribution of the species in between this site and the founder population is very scarce. Consistently, experimental data has revealed that *M. soledadinus* can survive flotation and exposures to saline conditions with half of the carabid beetles surviving flotation on seawater for ca. seven days (Renault, 2011; Hidalgo et al., 2013).

Large riversor rocky places without any coastal vegetation (e.g. cliffs) appeared to locally and temporarily reduce the spreading process of *M. soledadinus*. Conversely, dispersion can be fast along the seashore; assuming that flightless *M. soledadinus* invaded the south coast of Péninsule Courbet from the research station Port-aux-Français from 1996 onwards, the dispersal rate was estimated at 1.7 and 2.4 km / year eastward and westward, respectively. The comparison of the 1983 and 1995 observations on the north coast of Péninsule Courbet makes it possible to estimate this speed at 1.7 km/year. While Chevrier (1996) found a slightly higher dispersal speed of ca. 3.0 km / year on Ile Haute, the estimated dispersal rate of *M. soledadinus* was far lower in South Georgia (0.1 km / year, Brandjes et al., 1999) probably in connection with lower average temperatures on this island. Comparatively, the dispersal speed of a macropterous ground beetle – *Trechus obtusus* (body length: 3.2 – 4.3 mm) – was 3.0 km / year in Hawaii (Kavanaugh and Erwin, 1985; Liebherr and Takumi, 2002).

### 4.2. Mid-1990s, the arrival of M. soledadinus at the scientific research station Port-aux-Français

The arrival of *M. soledadinus* at Port-aux-Français (first observed in 1995) marked a significant milestone in the insect’s colonization of the archipelago in the sense that its displacement was subsequently assisted by human, as, for example, at Estacade. Estacade shelter has long been used for scientific purposes (monitoring of penguin populations) but also for tourists who overnighted at this location when visiting the penguin colonies. In 2005, the closest established populations of *M. soledadinus* were found at Cap Digby and at Port-aux-Français, more than 20 km far away from Estacade. The Estacade colonization likely resulted from the accidental transport of few individuals of *M. soledadinus* that were carried with food and material transported from Port-aux-Français. Similarly, the introduction of *M. soledadinus* to Port-Jeanne d’Arc, Port Phonolite (1991-1994) and Ile Guillou (1997) were most probably assisted by human activities.

Several islands of the Golfe du Morbihan, rarely visited, may have been colonized by specimens of *M. soledadinus* by flotation on algae (Renault, 2011; Hidalgo et al., 2013) and even through ornithochory. Kerguelen shags *Phalacrocorax verrucosus* use seawrecks that they sometimes bring from one island to another to build their nests. In addition, we found carrions and skulls of rabbits *Oryctolagus cuniculus* on the North-West coast of Ile Australia, whereas rabbit populations are absent. Carrions typically host *M. soledadinus* individuals which prey fly larvae (Renault et al. 2015); the transport and consumption of carrions by scavenging birds (i.e. skuas *Stercorarius antarcticus lonnbergi* and giant petrels *Macronestes giganteus*) thus represents another dissemination possibility of the insect.

During our survey (from December 2004 to March 2006), index 3 (> 100 individuals during a 10 min active search) and 2 (31-100 individuals) have been attributed 38 and 151 times respectively. On the west coast of Isthme Bas, we counted more than 150 individuals under a single stone (around 200 cm^2^). According to Chevrier (pers. comm.) such high abundances were never observed in the mid-1990s. Very high abundances (> 200 individuals during a 10 min active search) were recorded in 2013 at sites where *M. soledadinus* were moderately abundant (index 2, between Cap Cotter and Cap Digby, Port-aux-Français) or absent (east coast of Isthme Bas) in 2005-2006.

Abundances of *M. soledadinus* at Kerguelen Islands are much higher than those reported from South Georgia (maximum = 156 collected per hour, Brandjes et al., 1999) whereas it is now very easy to catch 4000 to 5000 specimens per hour in the surroundings of Port-aux-Français). Even though populations of *M. soledadinus* exhibit good abilities to cope with thermal stress (Engell Dahl et al. 2019, in press), the harsher climatic conditions encountered by the insect in South Georgia as compared with the Kerguelen Islands may reduce the speed of its geographic expansion. Mean annual temperature on King Edward Point during the period 1951-1980 was 2.0 °C and snow cover was more or less permanent from May to October in the coastal area (Ernsting 1993). Comparatively, the mean annual temperature is 4.4 °C at Port aux Français, and despite regular snowfalls (circa 15 days per month from June to August, Meteo France source), no permanent snow cover was observed over 1951-1980 at this site. Rising air temperature that occurred during winter months in the early 1990s may have amplified the spreading rate of *M. soledadinus* at Iles Kerguelen (Lebouvier et al., 2011), including at moderate altitudes (Lalouette et al., 2012; Laparie and Renault, 2016; Ouisse et al. 2019). The highest observation in the mid-1990s was 110m a.s.l. (Chevrier et al., 1997) and up to 358 m a.s.l. in 2005. In comparison, the highest observation inSouth Georgia was 275 m a.s.l. in 2009 (Convey et al., 2011). It should also be noted that in South Georgia glaciers can constitute impassable obstacles to dispersion.

### 4.3. Ecological impacts of the alien predatory M. soledadinus on native entomofauna

The invasion process of the predaceous ground beetle *M. soledadinus* strongly affects the native entomofauna even in the most recent colonised sites where specimens of *M. soledadinus* quickly became dominant. The native flies *A. aptera* and *C. moseleyi* seem to have totally disappeared in several locations colonised by *M. soledadinus* (Chevrier et al., 1997; Lebouvier et al., 2011, this study). When visiting Port-Couvreux in 2006, we found only a few specimens of *A. aptera* under seawrecks (one spot), down a small cliff at sea level (one spot) and under rabbit carrions (one spot at sea level and another at 288m a.s.l.). Furthermore, *C. moseleyi* appears to be more sensitive to *M. soledadinus* than *A. aptera* along seashore. *Merizodus soledadinus*, year-round active at the Kerguelen Islands (Ouisse et al. 2017), acts as a new ecological pressure which could now remove *C. moseleyi* from a secondary niche. Indeed, algae represent a secondary trophic niche for *C. moseleyi* whose primary resources are the Kerguelen cabbages (Tréhen et al., 1987) which almost disappeared from most locations invaded by rabbits (Chapuis et al., 1991). Further studies are underway to determine diet composition of *M. soledadinus* in habitats where both *A. aptera* and *C. moseleyi* locally became extinct.

Interactions of *M. soledadinus* are likely to occur with the three native arthropod predators: the rove beetle *Antarctophytosus atriceps* and the spiders *Myro kerguelensis* and *Neomaso antarcticus*. Direct predation may also occur; last-instar larvae and imagoes of *M. soledadinus* may eat small spiders and adults of *M. kerguelensis* could feed on larvae of the ground beetle. In South Georgia (Ernsting, 1993), 217 *Trechisibus antarcticus* and 68 spiders were found in one litter sample, all of them much smaller. Those spiders may well prey on springtails and be preyed upon by the carabid (Ernsting, 1993).

Although discreet at the first look as compared with the huge visible effect of the rabbit on plant communities with the significantly altered vegetation cover in the areas it colonized in the Kerguelen Islands (Chapuis et al., 1994), the impact of *M. soledadinus* on terrestrial ecosystems is potentially considerable. Despite its inability to fly, the distribution of the alien beetle in the archipelago is clearly expanding, and our results show that human activities may have contributed to this dispersion. If no control measures, let alone eradication, are practicable, it is essential to limit the transport of this invasive insect by human activities. This is why strict biosecurity measures (cleaning of clothing, shoes, equipment, and freight) have been put in place by the manager of the nature reserve for all travel on the Kerguelen Islands with even greater attention paid to visits to remote sites not yet colonized by *M. soledadinus*.

## Acknowledgments

This work was supported by the French Polar Institute (program IPEV 136 SUBANTECO); the CNRS (Zone Atelier Antarctique et Subantarctique); and the Agence Nationale de la Recherche (ANR-07-VULN-004, EVINCE). The authors are grateful to all the persons who helped collect data in the field.

## References

Allee, W.C., 1931. Animal aggregations, a study in general sociology. The University of Chicago Press, Chicago.

Alpert, P., Bone, E., Holzapfel, C., 2000. Invasiveness, invasibility and the role of environmental stress in the spread of non-native plants. Perspectives in Plant Ecology, Evolution and Systematics 3, 52–66. https://doi.org/10.1078/1433-8319-00004

Arnaud, P., Beurois, J., 1996. Les Armateurs du Rêve. Editions F. Jambois, Marseille.

Barbet-Massin, M., Rome, Q., Villemant, C., Courchamp, F., 2018. Can species distribution models really predict the expansion of invasive species? PLoS ONE 13, e0193085. https://doi.org/10.1371/journal.pone.0193085

Blackburn, T.M., Pyšek, P., Bacher, SI, Carlton, J.T., Duncan, R.P., Jarošik, V., Wilson, J.R.U., Richardson, D.M., 2011. A proposed unified framework for biological invasions. Trends in Ecology and Evolution 26, 333–339. https://doi.org/10.1016/j.tree.2011.03.023

Brandjes, G. J., Block, W., Ernsting, G. 1999. Spatial dynamics of two introduced species of carabid beetles on the sub-Antarctic island of South Georgia. Polar Biology 21, 326–334. https://doi.org/10.1007/s003000050369

Briot, C., 1990. Les frères Bossière : pionniers des Kerguelen. Recueil de l’Association des Amis du Vieux Havre 49, 113–143.

Brown, P. M. J., Thomas, C.E., Lombaert, E., Jeffries, D.L., Estoup, A., Lawson Handley, L.J., 2011. The global spread of *Harmonia axyridis* (Coleoptera: Coccinellidae): distribution, dispersal and routes of invasion. BioControl 56, 623–641. https://doi.org/10.1007/s10526-011-9379-1

Chabrerie, O., Massol, F., Facon, B., Thevenoux, R., Hess, M., Ulmer, R., Pantel, J.H., Braschi J., Amsellem, L., Baltora-Rosset, S., Tasiemski, A., Grandjean, F., Gibert, P., Chauvat, M., Affre, L., Thiébaut, G., Viard, F., Forey, E., Folcher, L., Boivin, T., Buisson, E., Richardson, D.M., Renault, D., 2019. Biological invasion theories: Merging perspectives from population, community and ecosystem scales. Preprints 2019, 2019100327, https://www.preprints.org/manuscript/201910.0327/v2

Carey, J.R., 1996. The incipient Mediterranean fruit fly population in California: implications for invasion biology. Ecology 77, 1690–1697. https://doi.org/10.2307/2265775

Casagranda, M.D., Roigt-Juňent, S., Szumik, C., 2009. Endemism at different spatial scales: an example with Carabidae (Coleoptera: Insecta) of austral South America. Revista Chilena de Historia Natural 82, 17–42. http://dx.doi.org/10.4067/S0716-078X2009000100002

Catford, J.A., Vest, P.A., Richardson, D.M., Pyšek, P., 2012. Quantifying levels of biological invasion: toward the objective classification of invaded and invasible ecosystems. Global Change Biology 18, 44–62. https://doi.org/10.1111/j.1365-2486.2011.02549.x

Chapuis, J.-L., Vernon, P., Frenot, Y., 1991. Fragilité des peuplements insulaires : exemple des îles Kerguelen, archipel subantarctique. in: Réactions des êtres vivants aux changements de l’environnement, PIREN, CNRS, pp. 235–248.

Chapuis, J.-L., Boussès, P., Barnaud, G., 1994. Alien mammals, impact and management in the French Subantarctic Islands. Biological Conservation 67, 97–104. https://doi.org/10.1016/0006-3207(94)90353-0

Chevrier, M., 1996. Introduction de deux espèces d’insectes aux îles Kerguelen : processus de colonisation et exemples d’interactions. Thèse, Université de Rennes 1, France.

Chevrier, M., Vernon, P., Frenot, Y., 1997. Potential effects of two alien insects on a sub-Antarctic wingless fly in the Kerguelen islands. in: Battaglia, B., Valancia, J., Walton D.W.H. (Eds.), Antarctic communities – Species, structure and survival, Cambridge University Press, Cambridge, pp. 424–431.

Convey, P., Key, R.S., Key, R.J.D., Belchier, M., Waller, C.L., 2011. Recent range expansion in non-native predatory beetles on sub-Antarctic South Georgia. Polar Biology 34, 597–602. https://doi.org/10.1007/s00300-010-0909-6

Costello, C. J., Solow, A. R., 2003. On the pattern of discovery of introduced species. Proceedings of the National Academy of Sciences USA 100, 3321–3323. https://doi.org/10.1073/pnas.0636536100

Crooks, J. A., 2005. Lag times and exotic species: The ecology and management of biological invasions in slow-motion. Ecoscience 12, 316–329. https://doi.org/10.2980/i1195-6860-12-3-316.1

Darlington, P. J., 1970. Coleoptera: Carabidae of South Georgia. Pacific Insects Monograph 23, 234.

David, P., Thébault, E., Anneville, O., Duyck, P.F., Chapuis, E., Loeuille, N., 2017. Impacts of invasive species on food webs: a review of empirical data. In Advances in Ecological Research vol. 56 – Networks of Invasion: A Synthesis of Concepts (ed. D. A. Bohan, A.J. Dumbrell and F. Massol), pp. 1–60. Academic Press.

Delépine, G., 2002. Histoires extraordinaires et inconnues dans les mers australes. Editions Ouest-France, Rennes.

Dreux, P., Galiana, D., Voisin, J.F., 1992. Acclimatation de *Merizodus soledadinus* Guérin dans l’archipel de Kerguelen (Coleoptera, Trechidae). Bulletin de la Société Entomologique de France 97, 219–221.

Engell, Dahl J., Bertrand, M., Pierre, A., Curtit, B., Pillard, C., Tasiemski, A., Convey, P., Renault, D., In Press. Thermal tolerance patterns of a carabid beetle sampled along invasion and altitudinal gradients at a sub-Antarctic island. Journal of Thermal Biology.

Enderlein, G., 1912. Die Insekten des Antarkto-Archiplata-Gebietes (Feuerland, Falklands-Inseln, Süd-Georgien). Konglica Svenska Vetenskapsakademiens Handlingar 48, 1–170.

Ernsting, G., 1993. Observations on life cycle and feeding ecology of two recently introduced predatory beetle species at South Georgia, sub-Antarctic. Polar Biology 13, 423–428. https://doi.org/10.1007/BF01681985

Frenot, Y., Gloaguen, J.C., Massé, L., Lebouvier, M., 2001. Human activities, ecosystem disturbance and plant invasions in subantarctic Crozet, Kerguelen and Amsterdam Islands. Biological Conservation 101, 33–50. https://doi.org/10.1016/S0006-3207(01)00052-0

Frenot, Y., Chown, S.L., Whinam, J., Selkirk, P.M., Convey, P., Skotnicki, M., Bergstrom, D.M., 2005. Biological invasions in the Antarctic: extent, impacts and implications. Biological Reviews 80, 45–72. https://doi.org/10.1017/S1464793104006542

Gurevitch, J., Padilla, D.K., 2004. Are invasive species a major cause of extinctions? Trends in Ecology and Evolution 19, 470–474. https://doi.org/10.1016/j.tree.2004.07.005

Hidalgo, K., Laparie, M., Bical, R., Larvor, V., Bouchereau, A., Siaussat, D., Renault, D., 2013. Metabolic fingerprinting of the responses to salinity in the invasive ground beetle *Merizodus soledadinus* at the Kerguelen Islands. Journal of Insect Physiology 59, 91–100. https://doi.org/10.1016/j.jinsphys.2012.10.017

Hui, C., Richardson, D.M., Landi, P., Minoarivelo, H.O., Garnas, J., Roy, H.E., 2016. Defining invasiveness and invasibility in ecological networks. Biological Invasions 18, 971.

Hullé, M., Buchard, C., Georges, R., Vernon, P., 2018. Guide d’identification des Invertébrés de Kerguelen et Crozet. 2nd édition. Université Rennes 1. https://doi.org/10.15454/1.5375302767618145E12

Hulme, P.E., Bacher, S., Kenis, M., Klotz S., Kühn, I., Minchin, D., Nentwig, W., Olenin, S., Panov, V., Pergl, J., Pyšek, P., Roques, A., Sol, D., Solarz, W., Vilà, M., 2008. Grasping at the routes of biological invasions: a framework for integrating pathways into policy. Journal of Applied Ecology 45, 403–414. https://doi.org/10.1111/j.1365-2664.2007.01442.x

Jeannel, R., 1940. Croisière du Bougainville aux Iles Australes françaises. Mémoires du Muséum National d’Histoire Naturelle, Paris 14, 63–201.

Jeannel, R., 1962. Les Tréchidés de la Paléantarctide occidentale. in: Delamare-Debouteville, C. and Rapoport, E. (Eds.), Biologie de l’Amérique Australe, Paris, pp. 527–655.

Jeannel, R., 1964. Biogéographie des Terres Australes de l’Océan Indien. Revue Française d’Entomologie 31, 319–417.

Jeschke, J. M., Strayer, D. L., 2005. Invasion success of vertebrates in Europe and North America. Proceedings of the National Academy of Sciences USA 102, 7198–7202. https://doi.org/10.1073/pnas.0504835102

Johns, P.M., 1974. Arthropoda of the subantarctic islands of New Zealand (1) Coleoptera : Carabidae Southern New Zealand, Patagonian, and Falkland Islands insular Carabidae. Journal of the Royal Society of New Zealand 4, 283–302. https://doi.org/10.1080/03036758.1974.10419396

Kadoya, T. Ishii, H.S., Kikuchi, R., Suda, S., Washitani, I. 2009. Using monitoring data gathered by volunteers to predict the potential distribution of the invasive alien bumblebee *Bombus terrestris*. Biological Conservation 142, 1011–1017. https://doi.org/10.1016/j.biocon.2009.01.012

Kavanaugh, D. H., Erwin T. L., 1985. *Trechus obtusus* Erichson (Coleoptera: Carabidae), a European ground beetle, on the Pacific coast of North America: its distribution, introduction, and spread. Pan-Pacific Entomologist 61, 170–179.

Kolar, C. S., Lodge, D. M., 2001. Progress in invasion biology: predicting invaders. Trends in Ecology and Evolution 16, 199–204. https://doi.org/10.1016/S0169-5347(01)02101-2

Lalouette, L., 2009. Impact de l’activité anthropique et des changements climatiques sur le succès envahissant de *Merizodus soledadinus* (Coleoptera, Carabidae) introduit aux Iles Kerguelen. Thèse de doctorat, Université de Lyon 1, 207 pp.

Lalouette, L., Williams, C.M., Cottin, M., Sinclair, B.J., Renault, D., 2012. Thermal biology of the alien ground beetle *Merizodus soledadinus* introduced to the Kerguelen Islands. Polar Biology 35, 509–517. https://doi.org/10.1007/s00300-011-1096-9

Laparie, M., Renault, D., 2016. Physiological responses to temperature in *Merizodus soledadinus* (Col., Carabidae), a subpolar carabid beetle invading sub-Antarctic islands. Polar Biology 39, 35–45. [https://doi.org/10.1007/s00300-014-1600-0]

Lebouvier, M., Laparie, M., Hullé, M., Marais, A., Cozic, Y., Lalouette, L., Vernon, P., Candresse, T., Frenot, Y., Renault, D., 2011. The significance of the sub-Antarctic Kerguelen Islands for the assessment of the vulnerability of native communities to climate change, alien insect invasions and plant viruses. Biological Invasions 13, 1195–1208. https://doi.org/10.1007/s10530-011-9946-5

Lefcheck, J.S., 2016. PIECEWISESEM: Piecewise structural equation modelling in R for ecology, evolution, and systematics 7, 573–579. https://doi.org/10.1111/2041-210X.12512

Levine, J.M., D’Antonio, C.M., 2003. Forecasting biological invasions with increasing international trade. Conservation Biology 17, 322–326.https://doi.org/10.1046/j.1523-1739.2003.02038.x

Liebherr, J. K., Takumi, R., 2002. Introduction and distributional expansion of *Trechus obtusus* (Coleoptera, Carabidae) in Maui, Hawai’i. Pacific Science 56, 365–375.

Lombaert, E., Guillemaud, T., Cornuet, J.M., Malausa, T., Facon, B., Estoup, A., 2010. Bridgehead effect in the worldwide invasion of the biocontrol Harlequin Ladybird. PLoS One 5, e9743. https://doi.org/10.1371/journal.pone.0009743

Lustig, A., Worner, S.P., Pitt, J.P., Doscher, C., Stouffer, D.B., Senay, S., 2017. A modeling framework for the establishment and spread of invasive species in heterogeneous environments. Ecology and Evolution 7, 8338–8348. https://doi.org/10.1002/ece3.2915

Niemelä, J., 1990. Habitat Distribution of Carabid Beetles in Tierra Del Fuego, South-America. Entomologica Fennica 1, 3–16. https://doi.org/10.33338/ef.83348

Ottesen, P. S., 1990. Diel activity patterns of Carabidae, Staphylinidae and Perimylopidae (Coleoptera) at South Georgia, sub-Antarctic. Polar Biology 10, 515–519. https://doi-org.passerelle.univ-rennes1.fr/10.1007/BF00233700

Ouisse, T., Laparie, M., Lebouvier, M., Renault D., 2017. New insights into the eco-biology of *Merizodus soledadinus*, a predatory carabid beetle invading the subantarctic Kerguelen Islands. Polar Biology 40, 2201–2209.

Ouisse, T., Day, E., Laville, L. Hendrickx, F., Convey P., Renault D., In Press, Does climate change facilitate the expansion of the invasive carabid beetle *Merizodus soledadinus* in the sub-Antarctic Kerguelen Islands? Scientific Reports.

Parker, I. M., Simberloff D., Lonsdale, W.M., Goodell, K., Wonham, M., Kareiva, P.M., Williamson, M.H., Von Holle, B., Moyle, P.B., Byers, J.E., Goldwasser, L., 1999. Impact: toward a framework for understanding the ecological effects of invaders. Biological Invasions 1, 3–19. https://doi.org/10.1023/A:1010034312781

Perrard, A., Haxaire, J., Rortais, A., Villemant, C., 2009. Observations on the colony activity of the Asian hornet *Vespa velutina* Lepeletier 1836 (Hymenoptera: Vespidae: Vespinae) in France. Annales de la Société Entomologique de France 45, 119–127. https://doi.org/10.1080/00379271.2009.10697595

Renault, D., 2011. Sea water transport and submersion tolerance as dispersal strategies for the invasive ground beetle *Merizodus soledadinus* (Carabidae). Polar Biology 34, 1591–1595. https://doi.org/10.1007/s00300-011-1020-3

Renault, D., Chevrier, M., Laparie, M., Vernon, P., Lebouvier M., 2015. Characterization of the microhabitats colonized by the alien ground beetle *Merizodus soledadinus* at the Kerguelen Islands. Revue d’Ecologie 70 (suppt 12), 28–32.

Renault, D., Laparie, M., McCauley, S.J., Bonte, D., 2018. Environmental adaptations, ecological filtering and dispersal, central to insect invasions. Annual Review of Entomology 63, 345–368.

Richardson, D.M., Pyšek, P., Carlton, J.T., 2011. A compendium of essential concepts and terminology in invasion ecology. in: Richardson, D.M. (Ed), Fifty Years of Invasion Ecology: The Legacy of Charles Elton, Wiley-Blackwell, Oxford, pp. 409–420.

Robinson, G. S., 1984. Insects of the Falkland Islands: a checklist and bibliography. Henry Ling Ltd., The Dorset Press, Dorchester.

Roy, H. E., Wajnberg, E. (Eds.), 2008. From biological control to invasion: the Ladybird Harmoniaaxyridis as a model species. Springer.

Schermann-Legionnet, A., Hennion, F., Vernon, P., Atlan, A., 2007. Breeding system of the subantarctic plant species *Pringleaantiscorbutica* R. Br. and search for potential insect pollinisators in the Kerguelen Islands. Polar Biology 30, 1183–1193. https://doi.org/10.1007/s00300-007-0275-1

Schliep, E.M., Lany, N.K., Zarnetske P.L., Schaeffer, R.N., Orians, C.M., Orwig, D.A., Preisser, E.L., 2018. Joint species distribution modelling for spatio-temporal occurrence and ordinal abundance. Global Ecology and Biogeography 27, 142–155. DOI: 10.1111/geb.12666

Simberloff, D., Rejmanek, M., 2011. Encyclopedia of biological invasions. University of California Press, Berkeley.

Sofaer, H.R., Jarnevich, C.S., Pearse, I.S., Smyth, R.L., Auer, S., Cook, G.L., Edwards, T.C., Guala, G.F., Howard, T.G., Morisette, J.T., Hamilton, H., 2019. Development and delivery of species distribution models to inform decision-making. BioScience 69, 544–557. https://doi.org/10.1093/biosci/biz045

Stevenson, M.D., Rossmo, D.K., Knell, R.K., Le Comber, S.C., 2012. Geographic profiling as a novel spatial tool for targeting the control of invasive species. Ecography 35, 704–715. https://doi.org/10.1111/j.1600-0587.2011.07292.x

Taylor, C.M., Hastings, A., 2005. Allee effects in biological invasions. Ecology Letters 8, 895–908. https://doi.org/10.1111/j.1461-0248.2005.00787.x

Thuiller, W., Richardson, D.M., Rouget, M., Procheş, Ş., Wilson, J.R.U., 2006. Interactions between environment, species traits, and human uses describe patterns of plant invasion. Ecology 87, 1755–1769. https://doi.org/10.1890/0012-9658(2006)87[1755:IBESTA]2.0.CO;2

Thuiller, W., Gassó, N.,Pino, J.,Vilà, M., 2012. Ecological niche and species traits: key drivers of regional plant invader assemblages. Biological Invasions 14, 1963–1980. https://doi.org/10.1007/s10530-012-0206-0

Tréhen, P., Voisin, J.F., 1984. Sur la présence de *Merizodussoledadinus* Guérin à Kerguelen (Coléoptère, Trechidae). L’Entomologiste 40, 53–54.

Tréhen, P., Vernon, P., Delettre, Y., Frenot, Y., 1987. Organisation et dynamique des peuplements diptérologiques à Kerguelen. Mise en évidence de modifications liées à l’insularité (exemple de l’Ile de Croÿ, Iles Nuageuses). Comité National Français des Recherches Antarctiques 58, 241–253.

Veldtman, R., Chown, S.L., McGeoch, M.A., 2010. Using scale-area curves to quantify the distribution, abundance and range expansion potential of an invasive species. Diversity and Distributions 16, 159–169. https://doi.org/10.1111/j.1472-4642.2009.00632.x

Vernon, P., 1981. Peuplement diptérologique des substrats enrichis en milieu insulaire subantarctique (Iles Crozet). Etude des Sphaeroceridae du genre Anatalanta. Université de Rennes I, Thèse Doctorat 3ème Cycle, 115 pp.

Voisin, J.-F., Chapelin-Viscardi, J.-D., Ponel, P., Rapp, M., 2017. Les Coléoptères de la province de Kerguelen (îles subantarctiques de l’océan Indien). Faune de France n°99. Fédération française des Sociétés de Sciences naturelles, Paris. 300 pp.

